# Fibromyalgia patients have altered lipid concentrations associated with disease symptom severity and anti-satellite glial cell IgG antibodies

**DOI:** 10.1101/2024.10.11.617524

**Authors:** Jenny E. Jakobsson, Joana Menezes, Emerson Krock, Matthew A. Hunt, Henrik Carlsson, Aina Vaivade, Payam Emami Khoonsari, Nilesh M. Agalave, Angelica Sandström, Diana Kadetoff, Jeanette Tour, Ida Erngren, Asma Al-Grety, Eva Freyhult, Katalin Sandor, Eva Kosek, Camilla I. Svensson, Kim Kultima

**Affiliations:** Department of Medical Sciences, Uppsala University, Uppsala, Sweden; Department of Surgical Sciences, Uppsala University, Uppsala, Sweden; Department of Physiology and Pharmacology, Center for Molecular Medicine, Karolinska Institutet, Stockholm, Sweden; Department of Clinical Neuroscience, Karolinska Institutet, Stockholm, Sweden; Department of Cell and Molecular Biology, Uppsala University, Uppsala, Sweden

**Keywords:** Fibromyalgia, Autoimmunity, Lipidomics, Lysophosphatidylcholine, Sphingomyelin

## Abstract

Autoimmunity and immunoglobulin G (IgG) autoantibodies may contribute to pain in a subset of fibromyalgia (FM) patients. Previously, we saw that IgG from FM patients induces pain-like behavior in mice and binds to satellite glial cells (anti-SGC IgG). The anti-SGC IgG levels were also associated with more severe symptomatology. Lipid metabolism in FM subjects is altered with lysophosphatidylcholines (LPCs) acting as pain mediators. The relationship between autoantibodies, lipid metabolism, and FM symptomatology remains unclear. We analyzed serum lipidomics with liquid chromatography mass spectrometry, anti-SGC IgG levels, and clinical measures in 35 female FM subjects and 33 age- and body mass index-balanced healthy controls (HC). Fibromyalgia subjects with higher anti-SGC IgG levels experienced more intense pain than those with lower levels. Sixty-three lipids were significantly altered between FM subjects and HC or between FM subjects with severe (FM severe) and mild symptoms (FM mild). Compared to HC, FM subjects had lower concentrations of lipid species belonging to the classes LPC (n = 10), lysophosphatidylethanolamine (n = 7), phosphatidylcholine (n = 4), and triglyceride (n = 5), but higher concentrations of diglyceride (n = 3). Additionally, FM severe had higher LPC 19:0, 22:0, and 24:1 and lower sphingomyelin (n = 9) concentrations compared to FM mild. A positive association was seen for LPC 22:0 and 24:1 with pain intensity and anti-SGC IgG levels in FM subjects. Taken together, our results suggest an association between altered lipid metabolism and autoimmune mechanisms in FM.

**Perspective:** Our results suggest an association between the postulated autoimmunity in FM and lipids that can act as pain mediators.

## Introduction

Fibromyalgia (FM) is a chronic pain syndrome characterized by widespread musculoskeletal pain, pain hypersensitivity, fatigue, sleep disturbances, depression, and anxiety^12,69^. It affects 2 - 4% of the general population, with a female:male ratio of 2:1^12,24,35^. Fibromyalgia pathogenesis is associated with generalized, multimodal increased sensitivity to cold and pressure stimuli as well as dysfunctional endogenous pain modulation^61^. Furthermore, neuroinflammation and widespread activation of cerebral glia have been reported in FM^1,3,36^. Although central sensitization is a well-established component of FM, increasing evidence suggests that peripheral mechanisms also contribute to the condition. Peripheral nerve pathology, in the form of reduced intraepidermal nerve fiber density (IENFD) and hyperexcitability of primary nociceptive afferents, is reported in approximately 50% of FM subjects^21^, which may lead to heightened pain perception and hypersensitivity to stimuli.

Systemic low-grade inflammation has been suggested to contribute to pain and other symptoms as elevated levels of certain pro- and anti-inflammatory cytokines have been reported^3,26,36^. Other forms of dysregulation of the immune system, including autoantibody production, have been suggested. In support of this notion, transferring immunoglobulin G (IgG) antibodies isolated from FM serum samples into mice elicits FM-like characteristics such as reduced IENFD, and increased nociceptor excitability^27^. In contrast, IgG from healthy controls (HC) had no discernible effect. Notably, IgG from individuals with FM exhibited a pronounced binding to satellite glial cells (SGCs) in both mouse and human dorsal root ganglia (DRGs). Furthermore, a higher degree of IgG binding to murine and human DRG SGCs was correlated to more severe FM symptoms in individual FM subjects^22,38^.

There is also evidence of altered muscle and lipid metabolism in FM, which has been linked to mitochondrial dysfunction and oxidative stress^15,25^. These metabolic abnormalities may contribute to impaired energy production and increased susceptibility to muscle fatigue and pain. Oxidative stress markers, such as byproducts of the oxidation of polyunsaturated fatty acids and phospholipids, have been related to the FIQ scores of FM subjects^60^. Furthermore, the lipidomic profile of FM plasma samples has shown differences in various lipid classes compared to healthy controls. Specifically, alterations have been observed in levels of lysophosphatidylcholine (LPC), lysophosphatidylethanolamine (LPE), phosphatidylcholine (PC), and sphingomyelin (SM)^6,30,46^. Dysregulation of lipid metabolism has been shown to influence the development of autoimmune diseases by affecting immune system functions and contributing to systemic inflammation, which is, for example, seen in rheumatic arthritis (RA) and systemic lupus erythematosus (SLE)^9^. Given the evidence that autoimmunity may also play a role in FM, it is also important to clarify the role of lipids in FM, as they could significantly influence the pathophysiology of FM.

Lipidomics provides a comprehensive view of lipid profiles, offering an understanding of complex biological systems and disease mechanisms, and insights that go beyond traditional targeted metabolite analyses. The untargeted approach further allows the identification of novel lipid species that may be previously unknown lipid mediators. In the present study, we utilized an untargeted lipidomics method to investigate lipid levelś variations in FM patients, who are part of a well-characterized cohort with an average of 9 years since diagnosis. Our objective was to explore how lipid variations correlate with FM symptoms, and the frequency of IgG binding to SGCs.

## Method and materials

### Participants

The current study is part of a larger project, with previously published genetic and neuroimaging results, as well as anti-SGC IgG binding dataset^19,20,22,23,56,66^. The study was approved by the regional ethical committee (2014/1604-31/1), and all participants signed written consent forms. Subjects, females aged 20-60 years, were recruited to Karolinska Institutet in Stockholm, Sweden, by advertising in the local press. FM subjects underwent a clinical examination by a pain specialist (Dr. Kadetoff), who also determined that they fulfilled the American College of Rheumatology (ACR) FM classification criteria from 1990^71^ and 2011^70^.

Exclusion criteria included having dominant pain conditions (other than FM), autoimmune or rheumatic diseases (other than FM), neurological disorders, other severe somatic diseases (such as cardiovascular and cancer), hypertension, pregnancy, body mass index (BMI) > 35 and smoking (> 5 cigarettes per day). Additional exclusion criteria were psychiatric disorders, including ongoing treatment for depression or anxiety, continuous medication with antidepressants or anticonvulsants, substance abuse, being unable to refrain from nonsteroidal anti-inflammatory drugs (NSAIDs), analgesics or hypnotics for at least 48 hours prior to study participation and not speaking Swedish. Healthy participants were screened by telephone to ensure they were free from all the exclusion criteria above and being free from chronic pain without regular medications with NSAIDs, analgesics, sleep medication, antidepressants, or anticonvulsants. The lipidomics data was only collected for individuals with BMI < 25 to limit confounding of body weight. After exclusion criteria were applied, 35 FM subjects and 33 HC remained.

### Questionnaires

Pain intensity was assessed with ratings on a 100 mm Visual Analogue Scale (VAS) where 0 represents no pain and 100 the worst imaginable pain. The ratings were noted for current (VASnow), minimum (VASmin), maximum (VASmax), and average (VASavr) pain intensity during the last week. Pain and FM duration (in months), total fibromyalgia impact questionnaire (FIQ) score, and number of tender points were noted for FM patients only. The FIQ assesses disease severity based on everyday life tasks, ranging from 0 to 100, where a higher score corresponds to a higher impairment^5^.

### Pressure algometry

Pressure pain thresholds (PPTs) were assessed with a handheld pressure algometer (Somedic Sales, AB) with a pressure rate increase corresponding to approximately 50 kPa/s and a probe area of one cm^2^. Assessments were made bilaterally at four different parts of the body: supraspinatus muscle, lateral epicondyle (elbow), gluteus muscle, and knee medial fat pad, once per site, and the average for each subject (PPTavr) was calculated and used for further analysis (described in detail by Fanton et al.^23^).

### Conditioned pain modulation

Conditioned pain modulation (CPM) was assessed using the Tourniquet test (ischemic pain, left forearm) as a conditioning stimulation. As described above, PPTs were assessed at the right thigh as the test stimulus. To induce ischemic pain, a blood pressure cuff (Tourniquet) was inflated to 200 mmHg, and the subjects were instructed to lift a one kg weight by extending their wrist until they rated the pain intensity as > 50 mm on a VAS, whereafter the forearm was resting and blood pressure cuff kept inflated. The PPTs were assessed twice before the tourniquet (baseline) and continuously during the tourniquet test (arm resting), with at least 10 second intervals between assessments and for a duration of four minutes or until the participant wished to end the test (PPTend). The CPM score was calculated as: each participant’s end PPT value − the first PPT baseline divided by the first PPT baseline (described in detail by Fanton et al.^23^).

### Blood sampling

After PPT and CPM assessment, blood was drawn from the median antecubital vein to BD vacutainer® STT II tubes (Fisher Scientific, 12917696). The collected blood was kept at room temperature for 40 minutes, followed by centrifugation at 2500 rpm for 10 minutes. The supernatant, comprising serum, was collected and aliquoted in Eppendorf tubes and stored at –80°C until further use.

### Primary SGCs culture

All performed procedures were approved by the local ethical committees (Stockholm Norra Djurförsöksetiska nämnd, 4945-2018). Adult female BALB/cAnNRj mice were euthanized using 5% isoflurane, followed by decapitation. Approximately 35 - 40 dorsal root ganglia (DRG) were harvested for primary cell culture, as previously described^27^. Succinctly, cells were dissociated with papain (Worthington, LS003126) and collagenase/dispase (Worthington, LS004176 and Sigma, D4693-1G) solutions for 30 minutes each with gentle shaking at 37°C. Later, cells were resuspended in 10% fetal bovine serum (FBS, Thermofisher, 10082147), 1x penicillin-streptomycin in F-12 media, then gently triturated and filtered in a 100 μm cell strainer (BD Bioscience, 352360) to reduce debris. The cell suspension was then added to non-coated Nunc™ Lab-Tek™ chamber slides (Thermofisher, 177445PK) for one and a half hours, and then the supernatant, which contained most neurons, was removed. Cells were recovered overnight at 37°C in a 5% CO2 incubator until serum incubation and immunocytochemistry the following day.

### Immunocytochemistry

The assessment of the frequency of serum IgG binding to SGCs (IgG+SGC%) was carried out as previously described^38^. In brief, SGCs were washed and incubated with individual patient or control serum previously filtered with a 0.22 μM filter and diluted 1:100 in culture media for three hours. By incubating the serum with living cells, IgG antibodies could only bind cell surface and extracellular antigens. After incubation, cells were washed, fixed for 10 minutes with 4% formaldehyde (PFA), and permeabilized for 5 minutes with 0.1% Triton-X100 in 1X PBS. Subsequently, cells were incubated with a rabbit glutamine synthetase (GS) IgG antibody (1:500, Abcam, ab73593), an SGC marker, overnight at 4°C. The next day, SGCs were washed and incubated with AF594 anti-human IgG antibody (1:300, Thermofisher, A11014) and AF488 anti-rabbit IgG antibody (1:300, Thermofisher, A11008) for one hour. After washing, cells were counterstained with Hoescht for 10 minutes. Slides were then left to dry for 10 minutes and coverslipped with Prolong gold mounting media. Cells were imaged with a Zeiss LSM800 confocal microscope.

### Image analysis

Images were analyzed using a custom pipeline based on the DRGquant pipeline^31^. In brief, a stardist^58^ model was trained to identify all Hoechst-stained nuclei, and a UNET^53^ model was trained to identify the soma of all cultured cells. The region of interest (ROI) of each nucleus was expanded to fill the soma of each cell. For identification of a positive signal for either GS (SGC marker) or IgG, 8xSD from all images’ pixel intensity average was used to set the minimum threshold of signal. Additionally, a summary image for each sample was generated in FIJI^57^, displaying all identified cells and their ROIs, which were visually inspected to assess and assure the quality of the analysis. Finally, a Python script was used to concatenate all data tables generated and format outputs that could easily be placed into GraphPad Prism (version 8.4.3, GraphPad Software, San Diego, California, USA). The generated IgG+SGC% dataset has previously been published by our group^22^.

### Untargeted lipidomic analysis

The serum samples were collected, prepared, and subjected to an untargeted lipidomics analysis using liquid chromatography coupled to high-resolution mass spectrometry (LC-HRMS) operating in polarity switching mode^8^, where positive and negative ionization was alternated during a single run.

#### Sample preparation

The methodology has previously been described in detail^8^. In brief, serum samples were thawed on ice, and 25 μL serum was precipitated using 75 μL of a precipitation solution made from isopropanol (IPA) supplemented with 14 isotopically labeled internal standards (IS) from Avanti Polar Lipids (Alabama, AL, USA). The IS included were cholesterol (d7), cholesterol ester (CE) (18:1-d7), diacylglycerol (diglyceride; DG) (15:0/18:1-d7), LPC (18:1-d7), monoacylglycerol (MG) (18:1-d7), PC (15:0/18:1-d7), phosphatidylserine (PS) (15:0/18:1-d7), phosphatidic acid (PA) (15:0/18:1-d7), triacylglycerol (triglyceride; TG) (15:0/18:1-d7/15:0), sphingomyelin (SM) (d18:1/18:1-d9), lysophosphatidylethanolamine (LPE) (18:1-d7), phosphatidylethanolamine (PE) (15:0/18:1-d7), phosphatidylinositol (PI) (15:0/18:1-d7) and phosphatidylglycerol (PG) (15:0/18:1-d7). The samples were left at –20°C for one hour before centrifugation at 14,800 rpm and 4°C. The supernatants were transferred to high-performance LC (HPLC) glass vials and stored at –80°C until analysis. Samples were prepared in batches containing up to 32 samples selected in a constrained randomized fashion to balance samples from patients and controls. Five µL of prepared sample in each batch was pooled to create a batch pool (QCB). Following the complete preparation of all batches, the QCB samples were pooled into a grand pool (QCG) containing all samples. The QCG sample was used as a quality control sample throughout the experiments. An external control (QCE) sample was prepared with all batches, along with blank samples containing water instead of serum prepared, with and without IS.

#### Mass spectrometry

In the LC-HRMS analysis, two µL of each sample was injected on a reversed-phase HPLC C18 column (Accucore C30 100 × 2.1 mm, 2.6 µm, Thermo Scientific, Waltham, MA, USA) using an Ultimate 3000 HPLC system (Thermo Scientific) interfaced to a high-resolution hybrid quadrupole Q Exactive Orbitrap MS (Thermo Scientific). A 15 minute long chromatographic program including a gradient was applied using the mobile phases 60:40 (v:v) H_2_O:methanol with 0.1% acetic acid (mobile phase A) and 90:10 (v:v) IPA:methanol with 0.1% acetic acid (mobile phase B). Samples were injected in a randomized batch order. Every batch was divided into three sample blocks. The three QC samples were injected between every sample block, and blanks were injected at the start and end of batches. The chromatography and HRMS settings have previously been described in detail^8^.

The HRMS analysis of all samples was performed in the full scan mode, collecting data in the profile mode with a resolution of 70,000 m/z. Polarity switching was used for these experiments, with the mass spectrometer switching between positive and negative polarity every second scan. We used a pooled sample containing aliquots of all samples to collect fragmentation data (MS/MS; MS2). The MS2 collection was performed for the positive and negative polarities separately. For this purpose, nitrogen evaporation concentrated the pooled sample about four times. The same pooled and concentrated sample was injected using seven overlapping m/z windows: 1) m/z 100 - 320; 2) m/z 318 - 428; 3) m/z 426 - 536; 4) m/z 534 - 644; 5) m/z 642 - 752; 6) m/z 750 - 860; 7) m/z 858 - 1200. For each m/z window, nine experiments with different collision energies were used (normalized collision energy, NCE): 10, 15, 20, 22, 25, 27, 30, 35, and 40. Apart from varying NCEs, the same settings were used for all MS2 experiments: resolution: 35,000, loop count: 10 (TopN), maximum injection time: 200 ms, and isolation window: 1.9 m/z, with data collected in the profile mode. The chromatography was done the same way as the main full scan experiments. This results in 63 MS2 experiments in positive and negative polarity, respectively.

### Statistical analysis

#### Immunoglobulin G binding to satellite glial cells

A k-means clustering based on VASavr, VASmax, and FIQ scores was performed to divide the FM patients, into those with severe (FM severe) and mild symptoms (FM mild), of our previously published IgG+SGC% dataset^22^. Twenty-five random starts were done with two centers, resulting in a division based on FM symptom severity. The IgG+SGC% was assessed with a Kruskal-Wallis test followed by Dunn’s post hoc test. Significant differences had a P value < 0.05. Data were analyzed using GraphPad Prism (version 8.4.3, GraphPad Software, San Diego, California USA).

#### Mass spectrometry data processing

The mass spectrometry (MS) data were split into positive and negative ionization and peak-picked with MSConvert^10^. FeatureFinder and FeatureLinker from OpenMS (version 2.3.0)^54^ were used in a Knime workflow^4^ to quantify and connect features across samples. The parameters used for FeatureFinder and FeatureLinker are previously described^8^, with the modification of a mass shift of ≤ 10 ppm allowed in positive and ≤ 7 ppm in negative mode and a retention time shift of 15 seconds. All further MS data analysis was performed using R Statistical Software (version 4.1.2)^51^. The intensities were log_2_ transformed and normalized with locally estimated scatterplot smoothing (loess) function from the R package limma^52^, to minimize the effect of weekly intensity decline.

Features with non-missing values (coverage) > 80% across FM and HC samples were subjected to identification using SIRIUS (version 4.9.12)^18^, the metaboigniter workflow^49^, the LIPID MAPS database^64^, and an in-house compound library. Only lipids from classes of the 14 isotopically labeled internal standards were considered. In SIRIUS and the metaboigniter workflow, a mass deviation < 8 ppm was allowed. A post-mass calibration was performed based on the putative IDs from the metaboigniter, SIRIUS, and compound library. The LIPID MAPS database was searched with the post-mass calibrated masses, allowing a ≤ 3 ppm mass deviation. Lipid abbreviations and classes are used according to the updated LIPID MAPS classification^42^.

Linear regression (LR) was used to assess the compound concentration differences, adjusting for age, BMI, and week of MS acquisition. The LR model was trained to assess differences between HC and FM subjects, as well as differences within the FM group. The LR was assessed with ANOVA F-tests, and the groups were compared with pairwise post hoc tests using t-statistics. The post hoc tests between HC, FM, FM mild, and FM severe subjects were calculated with the emmeans R package^41^. A difference in concentrations with a P < 0.05 was considered significant.

Spearman’s rank correlation was used to assess the association between compound concentrations and clinical parameters (pain duration, FM duration, tender points, VAS ratings of pain intensity, FIQ score, PPTavr, and CPM). The compound concentrations were corrected for age, BMI, and the week of MS acquisition. The IgG+SGC% was normally distributed, unlike the other clinical parameters, and was investigated for relationships with lipid concentrations with linear regression correcting for age, BMI, and week of MS acquisition.

## Results

### Fibromyalgia patients have elevated anti-SGC IgG levels

The FM patients were divided based on symptom severity (VASavr, VASmax, and FIQ scores) of the full cohort, using k-means clustering. The following subjects were part of the lipidomic analysis: FM mild, n = 14, and FM severe, n = 21 (Supplementary Figure S1). Subject characterization is presented in Table 1. Fibromyalgia subjects had significantly higher anti-SGC IgG levels (estimated by IgG+SGC%) than HC (P = 0.0045) (Figure 1). Of note, the IgG+SGC% dataset has been previously published^22^ and has been reused for our lipidomic study. Anti-SGC IgG levels of subjects from the full cohort can be found in Supplementary Figure S2. Fibromyalgia subjects with severe symptoms showed higher anti-SGC IgG levels compared to HC (P = 0.0042). Anti-SGC IgG levels were associated with higher pain intensity ratings of FM subjects (Figure 2). Therefore, these results indicate that FM subjects with higher anti-SGC IgG levels report higher pain intensity.

**Figure 1.**
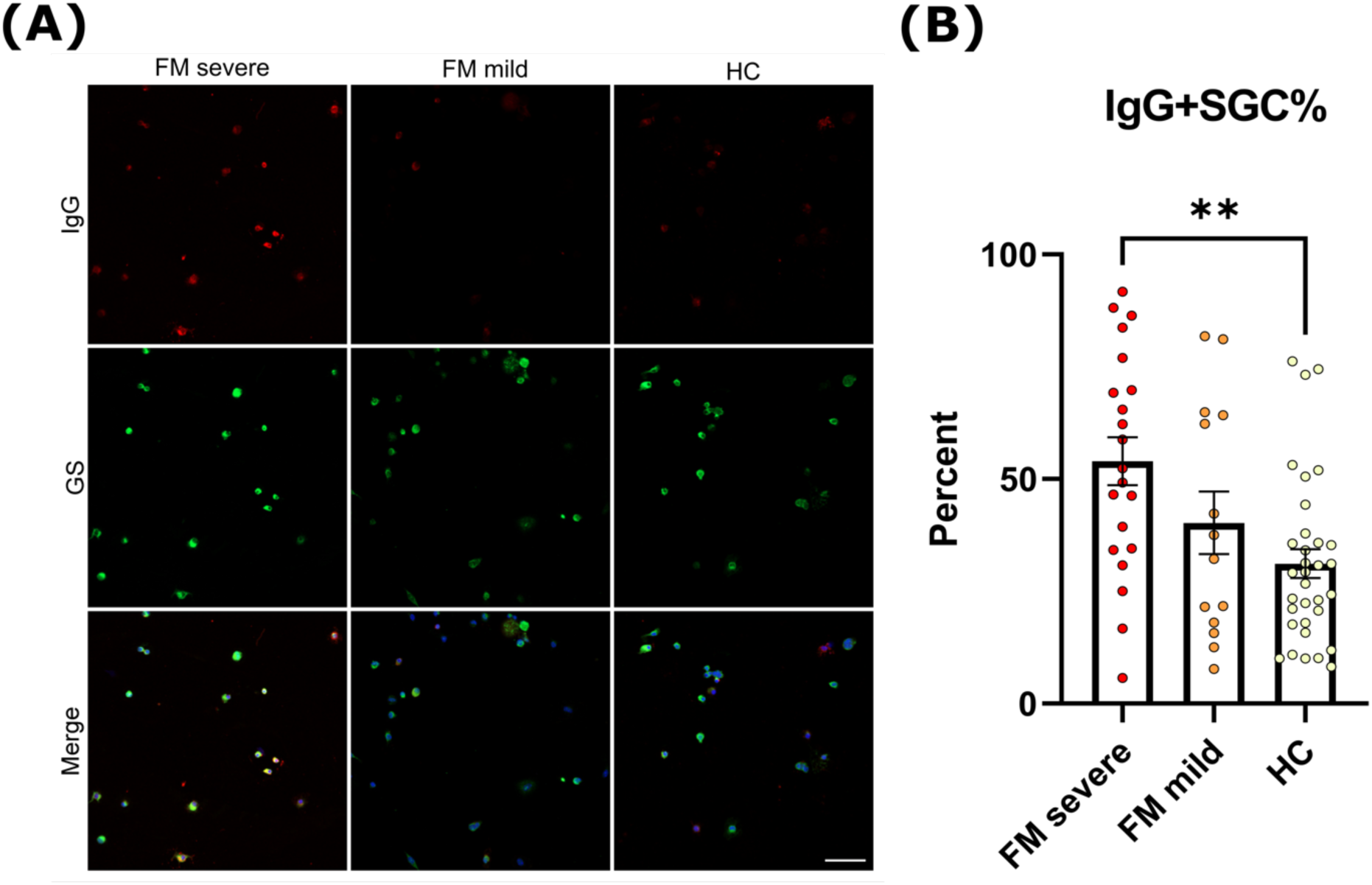
Representative images of satellite glial cell (SGC) cultures incubated with serum from fibromyalgia (FM) subjects with mild symptoms (FM mild) or severe symptoms (FM severe) and healthy controls (HC). The immunoglobulin G (IgG) and glutamine synthase (GS), an SGC marker, stainings shown are red and green, respectively. They are visualized separately and merged (A). The percentage of IgG-positive SGCs (IgG+SGC%) was compared between groups with a Kruskal-Wallis test followed by Dunn’s post hoc test (B). The scale bar is 50 μm. ***: P < 0.001, ****: P < 0.0001

**Figure 2.**
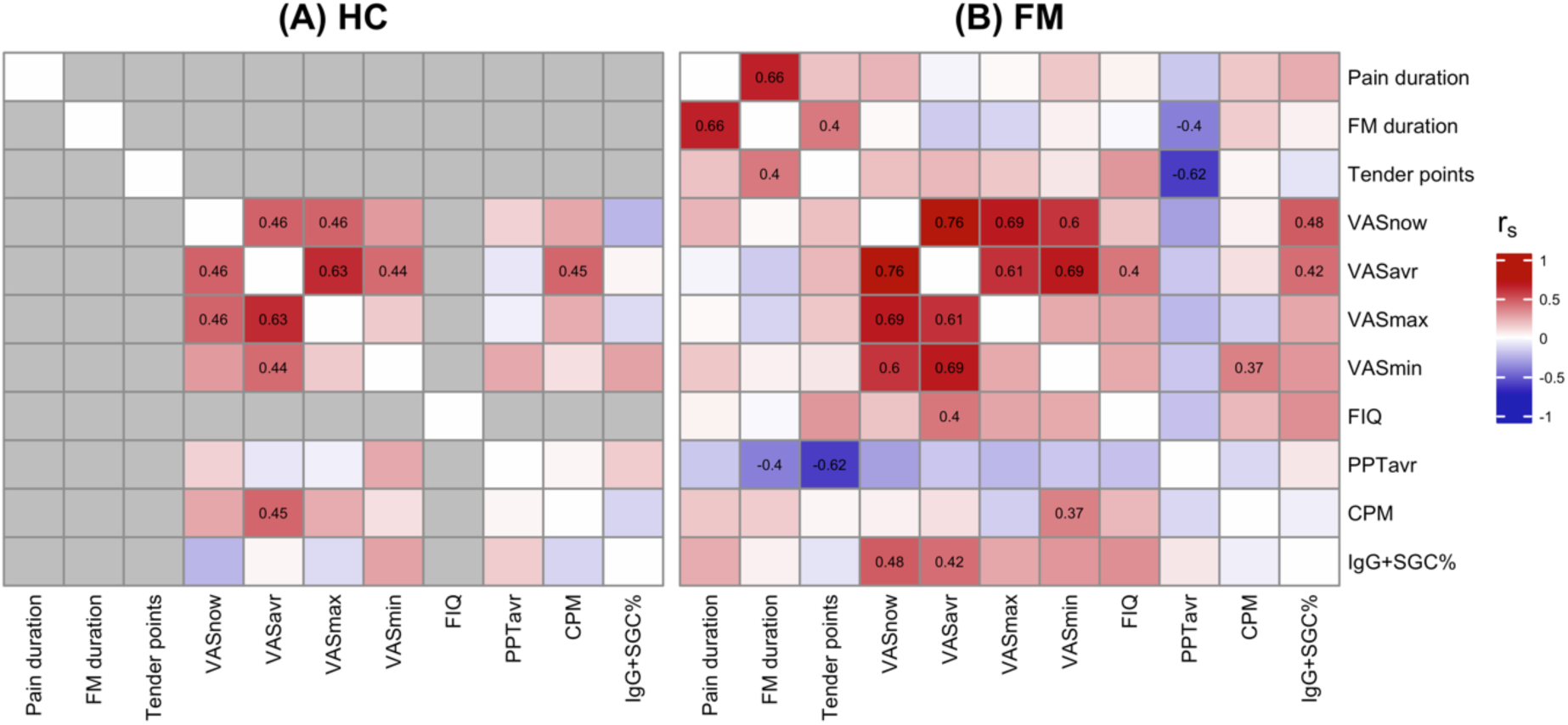
The frequency of immunoglobulin G binding to satellite glial cells (IgG+SGC%) in fibromyalgia (FM) subjects is associated with pain intensity ratings. The clinical parameters of healthy controls (HC; A) and FM subjects (B) were pairwise correlated to each other. The FM subjects had significant (P < 0.05) correlations between the frequency of immunoglobulin G binding to satellite glial cells (IgG+SGC%) and pain intensity ratings (Visual Analogue Scale; VAS). The numbers shown are Spearman’s correlation coefficients (r_s_) for the significant correlations (P < 0.05). The red color indicates a positive correlation, the blue a negative correlation, and the grey indicates that the data was not collected for that group.

**Table 1.** Subject characterization of the fibromyalgia (FM) subjects with mild (FM mild) and severe symptoms (FM severe) and healthy controls (HC). The values are present as mean ± standard deviation, except for the number of samples. The FM subjects were divided into subgroups: FM mild and FM severe. The tender points, FM duration, pain duration, and fibromyalgia impact questionnaire (FIQ) were only noted for the FM subjects. The pairwise group comparisons were performed with a Mann-Whitney U-test. NA: not applicable, *: P < 0.05

|  | HC | FM | FM mild | FM severe | FM VS HC | FM mild VS HC | FM severe VS HC | FM severe VS FM mild |
| --- | --- | --- | --- | --- | --- | --- | --- | --- |
| **Samples** | 33 | 35 | 14 | 21 | NA | NA | NA | NA |
| **Age** | 47.6±7.6 | 47.1±8.4 | 46.4±8.4 | 47.7±8.5 | P=0.8828 | P=0.5370 | P=0.8035 | P=0.4583 |
| **BMI** | 22.1±1.5 | 22.5±1.5 | 22.5±1.5 | 22.5±1.6 | P=0.2586 | P=0.5220 | P=0.2557 | P=1 |
| **Pain duration** | NA | 166.8±89.1 | 174.9±84 | 161.4±93.9 | NA | NA | NA | P=0.5777 |
| **FM duration** | NA | 117.2±78.5 | 125.1±79.1 | 111.9±79.6 | NA | NA | NA | P=0.5105 |
| **Tender points** | NA | 16.8±1.6 | 16.1±2 | 17.2±1.2 | NA | NA | NA | P=0.0732 |
| **VASnow** | 2±2.7 | 53.5±21.1 | 40.6±20.1 | 62.1±17.3 | P=1.897e-12* | P=6.913e-08* | P=6.056e-10* | P=0.0038* |
| **VASavr** | 4.7±6.3 | 57.1±18.2 | 41.1±14.5 | 67.8±11.2 | P=2.506e-12* | P=1.141e-07* | P=8.733e-10* | P=3.139e-05* |
| **VASmax** | 13.9±20.9 | 75.7±14.2 | 64.1±12.1 | 83.5±9.5 | P=5.840e-11* | P=1.997e-06* | P=4.396e-09* | P=5.229e-05* |
| **VASmin** | 2.6±4.9 | 33.1±21.8 | 23.6±16.2 | 39.3±23.1 | P=2.571e-11* | P=4.923e-07* | P=3.167e-09* | P=0.0570 |
| **FIQ** | NA | 61.4±14 | 51.9±10.5 | 67.8±12.5 | NA | NA | NA | P=0.0010* |
| **PPTavr** | 321.5±113.1 | 139.9±61.6 | 155.8±84.1 | 130.2±42.8 | P=7.569e-09* | P=0.0001* | P=1.203e-07* | P=0.4183 |
| **CPM** | 0.2±0.3 | -0.2±0.4 | -0.2±0.5 | -0.3±0.4 | P=2.112e-06* | P=0.0041* | P=5.577e-06* | P=0.8970 |
| **IgG+SGC%** | 31.2±18.5 | 48.5±25.6 | 40.2±26 | 53.9±24.5 | P=0.0045* | P=0.3625 | P=0.0006* | P=0.1105 |

### Lysophosphatidylcholine is the most commonly altered lipid class between FM subjects and HC

Linear regression was used to assess differences in compound concentration between FM (all, FM mild and FM severe) and HC. Out of 834 putatively identified lipids, we found 63 significantly altered lipids (ANOVA P < 0.05). All altered lipids are presented in Supplementary Table S1 and class-representative examples are shown in Figure 3 and Supplementary Figures S3-S9.

**Figure 3.**
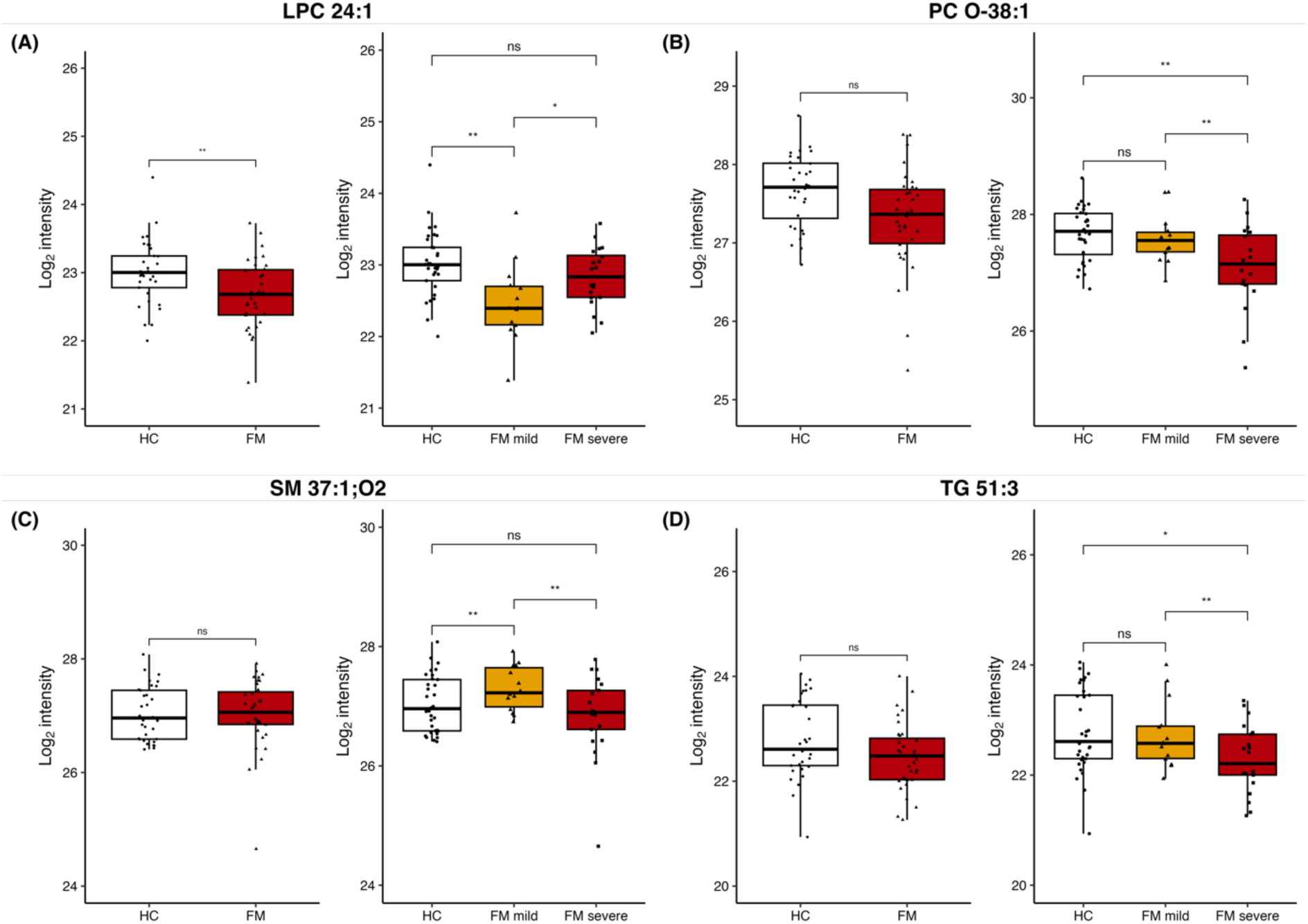
Lysophosphatidylcholines (LPCs), phosphatidylcholines (PCs), sphingomyelins (SMs), and triglycerides (TGs) are altered between fibromyalgia (FM) subjects and healthy controls (HC). The log_2_ mass spectrometry intensity signals (relative concentrations) of LPC 24:1 (A), PC O-38:1 (B), SM 37:1;O2 (C), and TG 51:3 (D) were stratified by FM subjects and HC or FM subjects with mild (FM mild) or severe symptoms (FM severe) and HC. ns: P ≥ 0.05, *: P < 0.05, **: P < 0.01

Fibromyalgia subjects had lower concentrations of LPCs (18:1, 18:2_1, 18:2_2, 18:3, 19:0, 20:2, 20:4, 20:5_1, 20:5_2, and 24:1), LPEs (16:0, 18:1, 18:3, 20:1, 20:2, 20:5, and 22:2), PCs (32:0, 34:0, 40:2 and 42:3), and TGs (54:1, 56:2, 56:4, 58:2, and 58:4) compared to HC (Supplementary Table S1). Higher concentrations of DGs (40:4_1, 40:5, and 42:6) were found in FM subjects compared to HC. The alkyl side chain lipid species did not follow the trend of their respective lipid classes, with DG O-32:0 being decreased and PC O-36:3 increased in FM subjects compared to HC. These results suggest that FM subjects typically have lower concentrations of LPCs, LPEs, PCs, and TGs than HC, whereas DGs are increased.

Furthermore, we investigated differences based on symptom severity by comparing FM severe and FM mild to HC and each other (Supplementary Table S1). Lysophosphatidylcholines (14:0, 18:1, 18:2_1, 18:2_2, 18:3, 20:2, 20:3, 20:4, 20:5_1, 20:5_2, 22:4, 22:5) and LPEs (16:0, 18:1, 18:3, 20:1, 20:2, 20:5, 22:2) had decreased concentrations in FM severe compared to HC. Although LPCs were found in lower concentrations in FM subjects and FM severe than in HC, LPCs 19:0, 22:0, and 24:1 were found in higher concentrations in FM severe compared to FM mild. Decreased concentrations of PCs were observed in FM severe (PC 32:0, 34:0, O-38:1, and O-38:4) and FM mild (PC 39:2, 40:2, 42:3, O-36:3, and O-42:1) compared to HC. Interestingly, there was again a divergence between alkyl or acyl side chains of PC species. Phosphatidylcholine 39:2 and 42:3 were found in higher concentrations in FM severe, while PC O-36:3, O-38:1, O-38:4, and O-42:1 were lower in FM severe compared to FM mild.

Sphingomyelins (SMs) (33:1;O2_2, 35:1;O2, 36:1;O2, 36:2;O3, 37:1;O2, and 39:2;O2) generally had increased concentrations in FM mild compared to HC, while generally non-changed between all FM subjects and HC (Supplementary Table S1). Additionally, SMs (32:1;O2_1, 32:1;O2_2, 32:1;O2_3, 33:1;O2_1, 33:1;O2_2, 35:1;O2, 37:1;O2, and 38:2;O2) were found in lower concentrations in FM severe than FM mild. TGs (50:2, 51:3, 51:5, 53:5, and 54:5_1) were also found in lower concentrations in FM severe compared to FM mild. These results suggest a difference in LPC, PC, TG, and SM concentrations depending on symptom severity in FM subjects.

### Lysophosphatidylcholine concentrations are associated with pain intensity ratings and anti-SGC IgG levels in FM subjects

To assess whether lipid concentrations were associated with FM symptom severity, we investigated the correlations to clinical parameters. Out of the 63 significantly (ANOVA P < 0.05) altered lipids, 22 correlated with pain intensity ratings independent of age and BMI (Figure 4). The alteration of the lipids with associations to pain is presented in Table 2. These pain-associated lipids were also investigated if they had an association with anti-SGC IgG levels (Table 2).

**Figure 4.**
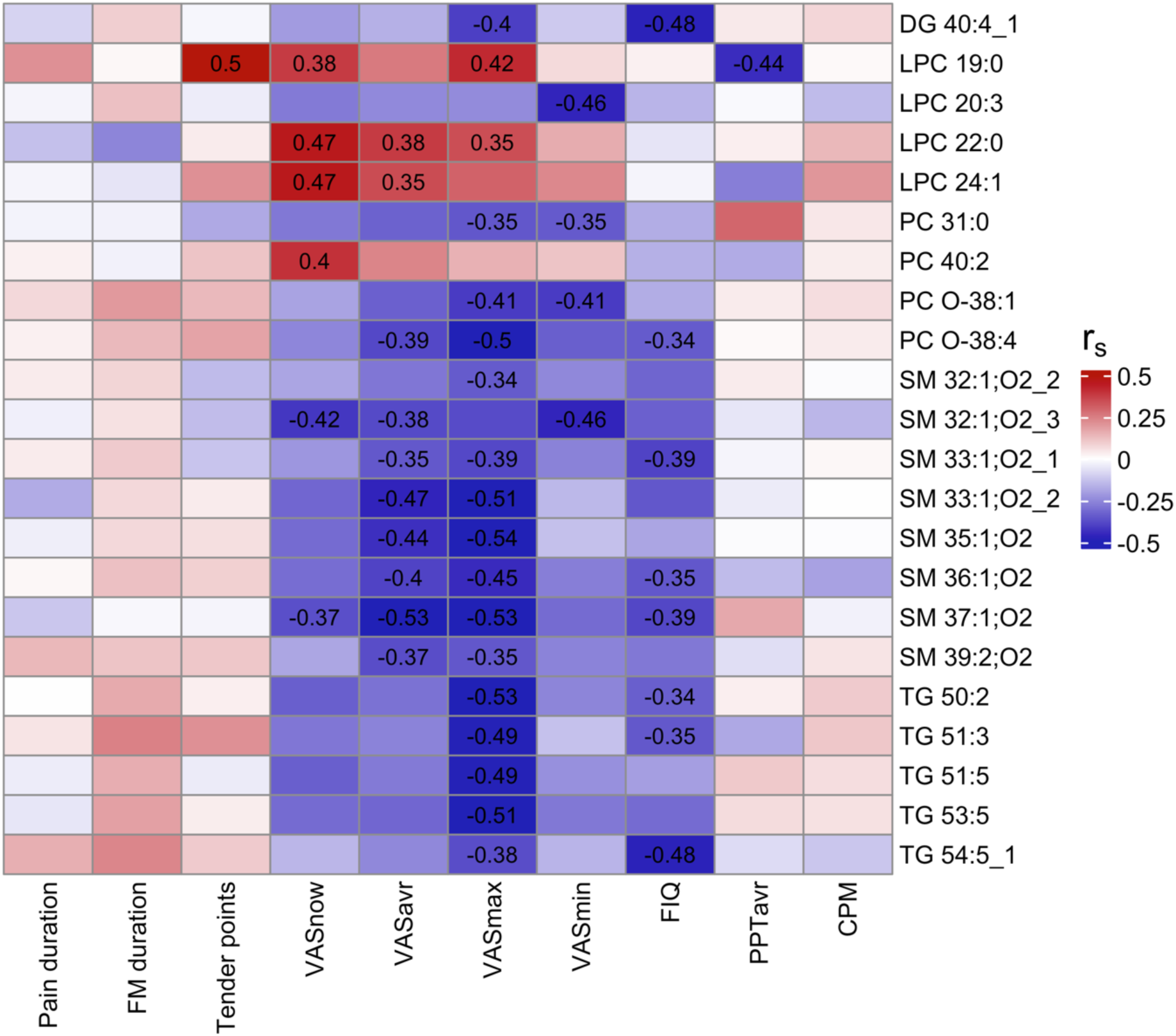
Lysophosphatidylcholines (LPCs) are positively correlated with pain intensity ratings, while the other lipid classes are negatively associated. The figure illustrates the correlation between the lipid concentrations and the demographic and clinical parameters in the fibromyalgia subjects (FMS). The majority of LPCs were found to positively correlate to pain intensity ratings (Visual Analogue Scale; VAS). In contrast, diglyceride (DG), phosphatidylcholines (PCs), sphingomyelins (SMs), and triglycerides (TGs) had negative associations with pain intensity ratings. The numbers shown represent Spearman’s rank correlation coefficients (r_s_) for the significant correlations (P < 0.05), adjusted for age, BMI and week of MS acquisition. A positive correlation is indicated by red, while a negative is indicated by blue.

**Table 2.** A summary of the significantly different lipids between fibromyalgia (FM) subjects with mild symptoms (FM mild) and severe symptoms (FM severe) and healthy controls (HC) with an association to pain intensity ratings. The log_2_ fold change (Est) was extracted from the linear regression models comparing HC and FM (FM mild and FM severe) or associations to IgG+SGC%, adjusting for age, BMI and week of MS acquisition. The table is sorted in alphabetical order by the abbreviated name. DG: diglyceride, LPC: lysophosphatidylcholine, PC: phosphatidylcholine, SM: sphingomyelin, TG: triglyceride

| Abbreviated<br>name | Name | ANOVA P | FM - HC |  | FM severe - HC |  | FM severe -<br>FM mild |  | FM mild - HC |  | IgG+SGC% |  |
| --- | --- | --- | --- | --- | --- | --- | --- | --- | --- | --- | --- | --- |
|  |  |  | Est | P | Est | P | Est | P | Est | P | Est | P |
| DG 40:4_1 | DG (22:2/18:2/0:0) | 0.0111 | 0.09 | 0.0085 | 0.05 | 0.2089 | -0.09 | 0.0674 | 0.13 | 0.0028 | -0.04 | 0.053 |
| LPC 19:0 | PC (19:0/0:0) | 0.0137 | -0.2 | 0.0468 | -0.03 | 0.764 | 0.34 | 0.0161 | -0.37 | 0.005 | 0.1 | 0.2323 |
| LPC 20:3 | PC (20:3/0:0) | 0.0389 | -0.15 | 0.2154 | -0.32 | 0.0204 | -0.35 | 0.0388 | 0.02 | 0.8787 | -0.12 | 0.2202 |
| LPC 22:0 | PC (22:0/0:0) | 0.0423 | -0.18 | 0.1187 | -0.01 | 0.9637 | 0.35 | 0.0309 | -0.36 | 0.018 | 0.15 | 0.0443 |
| LPC 24:1 | PC (24:1/0:0) | 0.0067 | -0.28 | 0.008 | -0.13 | 0.2769 | 0.3 | 0.0366 | -0.43 | 0.0016 | 0.14 | 0.0466 |
| PC 31:0 | PC (15:0/16:0) | 0.0493 | -0.1 | 0.5964 | -0.39 | 0.0636 | -0.58 | 0.0212 | 0.19 | 0.4059 | -0.11 | 0.4924 |
| PC 40:2 | PC (22:1/18:1) | 0.0407 | -0.36 | 0.0149 | -0.26 | 0.1167 | 0.2 | 0.3129 | -0.46 | 0.0155 | 0.07 | 0.5341 |
| PC O-38:1 | PC (O-18:0/20:1) | 0.0058 | -0.2 | 0.1075 | -0.43 | 0.0033 | -0.46 | 0.0087 | 0.03 | 0.8545 | -0.09 | 0.426 |
| PC O-38:4 | PC (O-18:0/20:4) | 0.0052 | -0.07 | 0.395 | -0.26 | 0.0102 | -0.36 | 0.0026 | 0.11 | 0.3179 | -0.09 | 0.1891 |
| SM 32:1;O2_2 | SM (d16:1/16:0) | 0.0264 | -0.04 | 0.6477 | -0.2 | 0.0489 | -0.31 | 0.0101 | 0.12 | 0.2936 | -0.06 | 0.4557 |
| SM 32:1;O2_3 | SM (d16:1/16:0) | 0.0279 | -0.01 | 0.9641 | -0.39 | 0.0904 | -0.76 | 0.0083 | 0.37 | 0.1453 | -0.26 | 0.1095 |
| SM 33:1;O2_1 | SM (d18:1/15:0) | 0.0109 | 0.01 | 0.9144 | -0.18 | 0.0795 | -0.38 | 0.0029 | 0.2 | 0.0866 | -0.09 | 0.2305 |
| SM 33:1;O2_2 | SM (d16:1/17:0) | 0.0386 | 0.27 | 0.2216 | -0.16 | 0.5221 | -0.87 | 0.0156 | 0.7 | 0.0233 | 0.4 | 0.0404 |
| SM 35:1;O2 | SM (d18:1/17:0) | 0.0131 | 0.05 | 0.6125 | -0.16 | 0.1678 | -0.43 | 0.0034 | 0.27 | 0.0474 | -0.08 | 0.355 |
| SM 36:1;O2 | SM (16:1;O2/20:0) | 0.0171 | 0.12 | 0.1385 | -0.03 | 0.7496 | -0.29 | 0.0085 | 0.26 | 0.0111 | -0.09 | 0.0981 |
| SM 37:1;O2 | SM(d18:1/19:0) | 0.0098 | 0.16 | 0.1477 | -0.06 | 0.6165 | -0.44 | 0.0043 | 0.38 | 0.0084 | -0.12 | 0.148 |
| SM 39:2;O2 | SM (d18:2/21:0) | 0.0142 | 0.21 | 0.0606 | 0.02 | 0.8982 | -0.39 | 0.0125 | 0.41 | 0.0058 | -0.13 | 0.1567 |
| TG 50:2 | TG (14:0/16:1/20:1) | 0.0469 | 0.03 | 0.6185 | -0.07 | 0.2802 | -0.19 | 0.0139 | 0.12 | 0.0869 | -0.02 | 0.735 |
| TG 51:3 | TG (12:0/17:2/22:1) | 0.005 | -0.02 | 0.8318 | -0.23 | 0.0311 | -0.42 | 0.0014 | 0.19 | 0.1115 | -0.03 | 0.6751 |
| TG 51:5 | TG (17:1/17:2/17:2) | 0.0482 | -0.12 | 0.5213 | -0.42 | 0.0534 | -0.6 | 0.023 | 0.18 | 0.467 | -0.02 | 0.8843 |
| TG 53:5 | TG (17:2/17:2/19:1) | 0.0346 | 0.05 | 0.7228 | -0.21 | 0.1991 | -0.51 | 0.0098 | 0.31 | 0.0942 | 0 | 0.9845 |
| TG 54:5_1 | TG (17:2/17:2/20:1) | 0.0404 | 0.12 | 0.1088 | 0.01 | 0.9313 | -0.22 | 0.0295 | 0.23 | 0.0169 | 0.01 | 0.8434 |

The concentrations of LPC 19:0, 22:0, and 24:1 were positively associated with pain intensity ratings (Figure 4). Furthermore, higher LPC 22:0 and 24:1 concentrations were also associated with higher anti-SGC IgG levels (Table 2). In conclusion, while LPCs were decreased in FM subjects compared to HC, they were positively related to pain intensity ratings and anti-SGC IgG levels in FM subjects.

In contrast, lipids belonging to the classes DG (40:4_1), PC (31:0, O-38:1, and O-38:4), SM (32:1;O2_2, 32:1;O2_3, 33:1;O2_1, 33:1;O2_2, 35:1;O2, 36:1;O2, 37:1;O2 and 39:2;O2) and TG (50:2, 51:3, 51:5, 53:5 and 54:5_1) were negatively correlated to pain intensity ratings. Additionally, DG 40:4_1, PC O-38:1, SM 33:1;O2_1, SM 36:1;O2, SM 37:1;O2, TG 50:2, TG 51:3, and TG 54:5_1 were negatively associated with FIQ scores and SM 33:1;O2_2 was associated with anti-SGC IgG levels. Thus, the decreased PC, SM, and TG concentrations in FM subjects, compared to HC, are also negatively associated with FM symptoms.

## Discussion

This study is the first to explore the relationship between lipids, clinical parameters, and autoimmunity in FM as measured in anti-SGC IgG levels. Our findings revealed significant alterations in the concentrations of lipid classes LPC, PC, SM, and TG when comparing serum from FM subjects to HC. Moreover, these lipid classes were also observed with associations with pain intensity ratings and anti-SGC IgG levels.

### The Role of LPC, LPE, and PC in FM

In our study, we observed decreased concentrations of PC species, such as PC O-38:1 and O-38:4, in the group with severe FM compared to mild FM and HC. Of note, we observed that the alteration of PC species varied depending on whether the side chain was acyl or alkyl. While the mechanistic implications of this are beyond the scope of the current study, this was a systemic trend observed in FM subjects.

LPCs have been shown to be pain mediators^34,37,50^. LPC can bind to and activate acid-sensing ion channel 3 (ASIC3), receptors typically proton-sensing and expressed in the peripheral nervous system^45^. The role of ASIC3 is well-established in inflammatory^16,44^ and joint pain^34,37^. Prior research has reported an increase in plasma LPC in FM patients, compared to HC, with some LPC species correlating with pain ratings in the FM group that experienced more severe and widespread pain^30^. Moreover, LPC 16:0 has been found to be increased in a stress-induced fibromyalgia-like animal model^30^. Our study found that the group with severe FM symptoms had higher LPC levels compared to those with milder. However, FM groups had lower LPC levels compared to HC. These differences might be attributed to variations in the time since diagnosis between studies; participants in the abovementioned study^30^ were recently diagnosed with FM, whereas the patients from our study had an average FM duration of nine years. Additionally, the previous study^30^ did not specify how the BMI was accounted for, whereas herein, we included only non-overweight FM patients. Decreased LPC concentrations have been observed in female adolescents with chronic pain^28,29,43^, chronic fatigue syndrome^28,29,43^, and persistent multisite musculoskeletal pain^28,29,43^, but no correlation to clinical parameters was performed in these studies. Herein, we observed a positive correlation between LPCs 19:0, 22:0, and 24:1 and pain intensity ratings. Importantly, higher LPC 22:0 and 24:1 concentrations were significantly associated with higher anti-SGC IgG levels in FM subjects, reinforcing a notable role of this lipid class in this disease.

Lysophosphatidylcholine species previously associated with neuropathic pain^39^, such as LPC 18:1 and 20:4, were found significantly decreased in FM subjects compared to HC, with no apparent correlation to clinical parameters in our study. In cerebrospinal fluid (CSF) from neuropathic pain patients, a lack of correlation of LPC with pain ratings was reported^39^. Interestingly, this study showed that lysophosphatidic acid (LPA), but not LPC and autotaxin (ATX) levels, significantly correlated to pain scores. Thus, LPC and ATX may not be directly linked to pain perception; instead, LPA appears to be a key pain mediator.

Lysophosphatidylethanolamines were found in lower concentrations in FM subjects compared to HC. Both LPC and LPE can serve as substrates for ATX^2^, an enzyme that synthesizes LPA. LPA plays a role in neuropathic pain^32,33^ and in murine models of FM^68^ and RA^63^. The decreased concentrations of LPC and LPE in FM subjects compared to HC may result from a faster turnover rate for these lipids, contributing to LPA production. Furthermore, LPC has suggested anti-inflammatory effects^59^, so lower levels of LPC may reflect a decreased immune suppression. This could potentially lead to the development of autoantibodies, such as those reported earlier^27^, through, e.g., reduced activation of G-protein-coupled receptor (GPCR) G2A receptors by LPC; which has been related to the development of autoimmunity due to alterations in T lymphocyteś function^17,40^.

### The Role of SM in FM

In FM patients, previous studies have reported an increase of SM 32:1;O2^46^, associated with musculoskeletal pain^48^. This is somewhat reflected in this study, where SMs were increased in FM mild compared to HC. However, we see decreasing levels of SMs in FM severe compared to FM mild, and a negative association with pain ratings and anti-SGC IgG levels, particularly SM 33:1;O2_2.

Sphingomyelin can be hydrolyzed by sphingomyelinases to ceramide, which can then be converted to sphingosine by ceramidases, and finally to sphingosine-1-phosphate (S1P) by the action of sphingosine kinases^62^. Plasma levels of S1P were previously reported to be higher in FM subjects compared to HC^46^, but associations with clinical parameters were not evaluated. In the current study, we did not directly quantify S1P levels, but the observed decreased SM levels may be due to increased SM hydrolysis to S1P. Elevated S1P levels could then increase neuronal and glia S1P receptor (S1PR) activation^62^, thereby exerting nociceptive and inflammatory effects. The significant linear relationship between anti-SGC IgG levels and SM 33:1; O2_2 shows that this specific lipid might participate in the feedback loop responsible for pain maintenance in FM after anti-SGC IgG is bound to SGC in the DRG, potentially promoting the release of pro-inflammatory mediators that could enhance SM 33:1; O2_2 production. However, the exact underlying mechanism needs to be further investigated. Therefore, future studies could further investigate SM metabolism and help elucidate its role in FM.

### The Role of TG in FM

Triglycerides tended to have lower concentrations in FM severe than in FM mild. They had negative correlations to vital clinical parameters, including VAS ratings and FIQ scores. Conversely, earlier studies have found an increase of serum TGs in FM patients compared to HC that correlated with clinical parameters, and that was hypothesized to be implicated in FM symptomatology in overweight patients^14,55^.

A recent study found decreased bile acid (BA) levels, specifically α-muricholic acid (α-MCA), in FM subjects compared to HC, which are concomitantly associated with clinical factors^47^. This is of interest since BAs are synthesized from cholesterol and have been correlated to serum triglycerides^11^. Our findings could point towards changes in BA metabolism and exacerbation of the disease. This would agree with the aforementioned study that points towards changes in BAs present in the serum of FM patients. Bile acid metabolism can be influenced by gut microbiota^7^. Interestingly, recent studies have reported a distinct microbiome in FM patients^7,13^ This alteration of gut species could result in an immune response that triggers FM. This might explain the elevated anti-SGC IgG observed and aid in the quest for antigen identification. However, this hypothesis remains highly speculative, and further studies, including longitudinal ones, are warranted to establish a definitive link. In our study, no significant predictions between TGs and anti-SGC IgG levels could be found.

In this study no strong associations of QST measures, such as PPT and CPM, with lipid concentrations and anti-SGC IgG levels were found. Therefore, we reason that alterations in lipid metabolism and the autoantibodies reported earlier may play an important role in spontaneous, rather than evoked pain in FM.

### Limitations

In this study, no causal claims are reported, solely associations. Furthermore, subjects were not asked to fast overnight, which can alter serum lipid levels. A limitation to our conclusions is that LPA levels were not analyzed in this study, due to inherent limitations associated with the LC-HRMS method used^65,67^. Finally, we have a limited number of subjects, which would ideally be validated in a larger cohort.

## Conclusion

Our findings reveal significant alterations in lipid concentration, especially for lipid classes PC, LPC, SM, and TG, when comparing serum samples from FM subjects with severe and mild symptoms and HC. Lysophosphatidylcholines were positively associated with pain intensity while remaining lipids were negatively associated. The lower concentrations of LPC and LPE in FM can be due to the conversion into LPA, and act pro-nociceptive. Concomitantly, SM might also be converting to S1P and playing a potential role in pain in FM. Furthermore, the association between LPCs and anti-SGC antibodies suggests an interaction between altered lipid metabolism and the autoimmune component previously described.

