## Supplementary material for "Fibromyalgia patients have altered lipid concentrations associated with disease symptom severity and anti-satellite glial cell IgG antibodies"

#### Supplementary Tables

| Abbreviated name | Matched name | ANOVA P | FM - HC | | FM severe - HC | | FM severe - FM mild | | FM mild - HC | |
|---|---|---|---|---|---|---|---|---|---|---|
| | | | Est | P | Est | P | Est | P | Est | P |
| **DG 40:4_1** | DG (22:2/18:2/0:0) | 0.0111 | 0.09 | 0.0085 | 0.05 | 0.2089 | -0.09 | 0.0674 | 0.13 | 0.0028 |
| **DG 40:4_2** | DG (18:2/22:2/0:0) | 0.0272 | -0.01 | 0.951 | -0.23 | 0.093 | -0.45 | 0.0082 | 0.22 | 0.1637 |
| **DG 40:5** | DG (18:2/22:3/0:0) | 0.0249 | 0.15 | 0.0131 | 0.09 | 0.1805 | -0.12 | 0.1439 | 0.21 | 0.0073 |
| **DG 42:6** | DG (20:3/22:3/0:0) | 0.0191 | 0.27 | 0.011 | 0.16 | 0.1833 | -0.22 | 0.116 | 0.38 | 0.0053 |
| **DG O-32:0** | DG (O-16:0/16:0) | 0.0162 | -0.46 | 0.0051 | -0.49 | 0.0088 | -0.06 | 0.7781 | -0.43 | 0.0378 |
| **LPC 14:0** | PC (14:0/0:0) | 0.0375 | -0.18 | 0.2063 | -0.38 | 0.019 | -0.4 | 0.0422 | 0.02 | 0.9093 |
| **LPC 18:1** | PC (18:1/0:0) | 0.0287 | -0.25 | 0.0307 | -0.35 | 0.0079 | -0.2 | 0.1933 | -0.15 | 0.313 |
| **LPC 18:2_1** | PC (18:2/0:0) | 0.0033 | -0.45 | 0.0034 | -0.58 | 0.0008 | -0.28 | 0.171 | -0.31 | 0.1087 |
| **LPC 18:2_2** | PC (18:2/0:0) | 0.0186 | -0.4 | 0.0204 | -0.56 | 0.005 | -0.32 | 0.1663 | -0.24 | 0.2577 |
| **LPC 18:2_3** | PC (18:2/0:0) | 0.0173 | 0.49 | 0.0052 | 0.5 | 0.0107 | 0.02 | 0.9407 | 0.48 | 0.0303 |

| | | | | | | | | | | |
|---|---|---|---|---|---|---|---|---|---|---|
| **LPC 18:3** | PC (18:3/0:0) | 0.0224 | -0.43 | 0.0232 | -0.6 | 0.0061 | -0.33 | 0.1954 | -0.26 | 0.2707 |
| **LPC 19:0** | PC (19:0/0:0) | 0.0137 | -0.2 | 0.0468 | -0.03 | 0.764 | 0.34 | 0.0161 | -0.37 | 0.005 |
| **LPC 20:2** | PC (20:2/0:0) | 0.0306 | -0.36 | 0.0117 | -0.4 | 0.0131 | -0.08 | 0.6621 | -0.32 | 0.079 |
| **LPC 20:3** | PC (20:3/0:0) | 0.0389 | -0.15 | 0.2154 | -0.32 | 0.0204 | -0.35 | 0.0388 | 0.02 | 0.8787 |
| **LPC 20:4** | PC (20:4/0:0) | 0.0012 | -0.26 | 0.0133 | -0.44 | 0.0004 | -0.36 | 0.0134 | -0.08 | 0.5353 |
| **LPC 20:5_1** | PC (20:5/0:0) | 0.0212 | -0.57 | 0.0075 | -0.63 | 0.0097 | -0.11 | 0.7007 | -0.52 | 0.0567 |
| **LPC 20:5_2** | PC (20:5/0:0) | 0.0134 | -0.5 | 0.0052 | -0.55 | 0.0061 | -0.1 | 0.6842 | -0.45 | 0.0522 |
| **LPC 22:0** | PC (22:0/0:0) | 0.0423 | -0.18 | 0.1187 | -0.01 | 0.9637 | 0.35 | 0.0309 | -0.36 | 0.018 |
| **LPC 22:4** | PC (22:4/0:0) | 0.0357 | -0.2 | 0.1248 | -0.37 | 0.0138 | -0.34 | 0.0592 | -0.03 | 0.8543 |
| **LPC 22:5** | PC (22:5/0:0) | 0.0151 | -0.22 | 0.0585 | -0.38 | 0.005 | -0.31 | 0.0514 | -0.07 | 0.6492 |
| **LPC 24:1** | PC (24:1/0:0) | 0.0067 | -0.28 | 0.008 | -0.13 | 0.2769 | 0.3 | 0.0366 | -0.43 | 0.0016 |
| **LPE 16:0** | PE (16:0/0:0) | 0.048 | -0.28 | 0.0231 | -0.33 | 0.0171 | -0.11 | 0.5139 | -0.23 | 0.1489 |
| **LPE 18:1** | PE (18:1/0:0) | 0.0045 | -0.53 | 0.0043 | -0.69 | 0.0011 | -0.33 | 0.1873 | -0.37 | 0.1161 |
| **LPE 18:3** | PE (18:3/0:0) | 0.0467 | -0.5 | 0.0301 | -0.65 | 0.0145 | -0.29 | 0.3588 | -0.36 | 0.2225 |
| **LPE 20:1** | PE (20:1/0:0) | 0.034 | -0.24 | 0.0174 | -0.29 | 0.0115 | -0.1 | 0.4583 | -0.19 | 0.1381 |
| **LPE 20:2** | PE (20:2/0:0) | 0.0103 | -0.35 | 0.0049 | -0.41 | 0.0036 | -0.11 | 0.5106 | -0.3 | 0.0621 |

| | | | | | | | | | | |
|---|---|---|---|---|---|---|---|---|---|---|
| **LPE 20:5** | PE (20:5/0:0) | 0.006 | -0.6 | 0.0021 | -0.67 | 0.0028 | -0.12 | 0.6308 | -0.54 | 0.029 |
| **LPE 22:2** | PE (22:2/0:0) | 0.0093 | -0.34 | 0.0075 | -0.44 | 0.0025 | -0.2 | 0.2431 | -0.24 | 0.133 |
| **PC 31:0** | PC (15:0/16:0) | 0.0493 | -0.1 | 0.5964 | -0.39 | 0.0636 | -0.58 | 0.0212 | 0.19 | 0.4059 |
| **PC 32:0** | PC (17:0/15:0) | 0.0097 | -0.14 | 0.0154 | -0.2 | 0.0024 | -0.12 | 0.1143 | -0.08 | 0.2836 |
| **PC 34:0** | PC (19:0/15:0) | 0.0239 | -0.14 | 0.0331 | -0.2 | 0.0065 | -0.13 | 0.1327 | -0.07 | 0.3834 |
| **PC 39:2** | PC (19:1/20:1) | 0.0367 | -0.17 | 0.3555 | 0.15 | 0.4633 | 0.64 | 0.0132 | -0.49 | 0.0372 |
| **PC 40:2** | PC (22:1/18:1) | 0.0407 | -0.36 | 0.0149 | -0.26 | 0.1167 | 0.2 | 0.3129 | -0.46 | 0.0155 |
| **PC 42:3** | PC (18:3/24:0) | 0.0015 | -0.24 | 0.0111 | -0.05 | 0.6548 | 0.39 | 0.0038 | -0.43 | 0.0005 |
| **PC O-36:3** | PC (O-16:0/20:3) | 0.0148 | 0.27 | 0.0467 | 0.05 | 0.745 | -0.44 | 0.0162 | 0.49 | 0.0054 |
| **PC O-38:1** | PC (O-18:0/20:1) | 0.0058 | -0.2 | 0.1075 | -0.43 | 0.0033 | -0.46 | 0.0087 | 0.03 | 0.8545 |
| **PC O-38:4** | PC (O-18:0/20:4) | 0.0052 | -0.07 | 0.395 | -0.26 | 0.0102 | -0.36 | 0.0026 | 0.11 | 0.3179 |
| **PC O-42:1** | PC (P-20:0/22:0) | 0.0373 | 0.17 | 0.2642 | -0.09 | 0.5957 | -0.52 | 0.014 | 0.43 | 0.0319 |
| **PE 38:2** | PE (P-18:0/20:1) | 0.0142 | -0.13 | 0.0263 | -0.19 | 0.0037 | -0.13 | 0.101 | -0.06 | 0.3813 |
| **PE O-36:2** | PE (P-18:0/18:1) | 0.0403 | 0.08 | 0.5367 | -0.15 | 0.3133 | -0.46 | 0.0121 | 0.31 | 0.0671 |
| **PE O-40:5** | PE (P-18:0/22:4) | 0.0288 | -0.07 | 0.58 | -0.28 | 0.0441 | -0.43 | 0.012 | 0.15 | 0.3467 |
| **SM 32:1;O2_1** | SM (d18:1/14:0) | 0.0485 | -0.03 | 0.8071 | -0.21 | 0.0941 | -0.37 | 0.0169 | 0.16 | 0.2674 |

| | | | | | | | | | | |
|---|---|---|---|---|---|---|---|---|---|---|
| **SM 32:1;O2_2** | SM (d16:1/16:0) | 0.0264 | -0.04 | 0.6477 | -0.2 | 0.0489 | -0.31 | 0.0101 | 0.12 | 0.2936 |
| **SM 32:1;O2_3** | SM (d16:1/16:0) | 0.0279 | -0.01 | 0.9641 | -0.39 | 0.0904 | -0.76 | 0.0083 | 0.37 | 0.1453 |
| **SM 33:1;O2_1** | SM (d18:1/15:0) | 0.0109 | 0.01 | 0.9144 | -0.18 | 0.0795 | -0.38 | 0.0029 | 0.2 | 0.0866 |
| **SM 33:1;O2_2** | SM (d16:1/17:0) | 0.0386 | 0.27 | 0.2216 | -0.16 | 0.5221 | -0.87 | 0.0156 | 0.7 | 0.0233 |
| **SM 35:1;O2** | SM (d18:1/17:0) | 0.0131 | 0.05 | 0.6125 | -0.16 | 0.1678 | -0.43 | 0.0034 | 0.27 | 0.0474 |
| **SM 36:1;O2** | SM (d16:1/20:0) | 0.0171 | 0.12 | 0.1385 | -0.03 | 0.7496 | -0.29 | 0.0085 | 0.26 | 0.0111 |
| **SM 36:2;O2** | SM (d18:1/18:1) | 0.031 | 0.2 | 0.0353 | 0.08 | 0.466 | -0.25 | 0.0592 | 0.32 | 0.009 |
| **SM 37:1;O2** | SM (d18:1/19:0) | 0.0098 | 0.16 | 0.1477 | -0.06 | 0.6165 | -0.44 | 0.0043 | 0.38 | 0.0084 |
| **SM 39:2;O2** | SM (d18:2/21:0) | 0.0142 | 0.21 | 0.0606 | 0.02 | 0.8982 | -0.39 | 0.0125 | 0.41 | 0.0058 |
| **TG 50:2** | TG (14:0/16:1/20:1) | 0.0469 | 0.03 | 0.6185 | -0.07 | 0.2802 | -0.19 | 0.0139 | 0.12 | 0.0869 |
| **TG 51:3** | TG (12:0/17:2/22:1) | 0.005 | -0.02 | 0.8318 | -0.23 | 0.0311 | -0.42 | 0.0014 | 0.19 | 0.1115 |
| **TG 51:5** | TG (17:1/17:2/17:2) | 0.0482 | -0.12 | 0.5213 | -0.42 | 0.0534 | -0.6 | 0.023 | 0.18 | 0.467 |
| **TG 52:5** | TG (16:1/18:2/18:2) | 0.0433 | 0.13 | 0.4104 | 0.37 | 0.0382 | 0.49 | 0.0247 | -0.12 | 0.564 |
| **TG 53:5** | TG (17:2/17:2/19:1) | 0.0346 | 0.05 | 0.7228 | -0.21 | 0.1991 | -0.51 | 0.0098 | 0.31 | 0.0942 |

| | | | | | | | | | | |
|---|---|---|---|---|---|---|---|---|---|---|
| **TG 54:1** | TG (18:0/18:0/18:1) | 0.0027 | -0.6 | 0.0007 | -0.6 | 0.0025 | 0 | 0.9858 | -0.61 | 0.0075 |
| **TG 54:5_1** | TG (17:2/17:2/20:1) | 0.0404 | 0.12 | 0.1088 | 0.01 | 0.9313 | -0.22 | 0.0295 | 0.23 | 0.0169 |
| **TG 54:5_2** | TG (18:1/18:2/18:2) | 0.01 | 0.08 | 0.6309 | 0.41 | 0.0279 | 0.67 | 0.0035 | -0.26 | 0.2102 |
| **TG 56:2** | TG (12:0/22:1/22:1) | 0.0053 | -0.67 | 0.0013 | -0.6 | 0.0096 | 0.14 | 0.6087 | -0.74 | 0.005 |
| **TG 56:4** | TG (17:1/17:2/22:1) | 0.0395 | -0.35 | 0.0116 | -0.33 | 0.0312 | 0.03 | 0.8849 | -0.36 | 0.0397 |
| **TG 58:2** | TG (17:1/19:0/22:1) | 0.0215 | -0.57 | 0.0121 | -0.34 | 0.1766 | 0.45 | 0.1348 | -0.8 | 0.0061 |
| **TG 58:4** | TG (17:2/19:1/22:1) | 0.0336 | -0.48 | 0.0096 | -0.42 | 0.0449 | 0.14 | 0.5789 | -0.55 | 0.0219 |

**Supplementary Table S1. Summary of the lipids found altered between fibromyalgia (FM) subjects and healthy control (HC).** There were 63 lipids found significantly ($P < 0.05$) altered between the groups. The $\log_2$ fold change (Est) was extracted from the linear regression models comparing HC and FM subjects, FM subjects with mild symptoms (FM mild), and FM subjects with severe symptoms (FM severe). The most common lipid class found was lysophosphatidylcholine (LPC), followed by triglyceride (TG), sphingomyelin (SM), and phosphocholines (PC). There are duplicated lipids, but these either have different adducts or retention times (which may result from different positions of the double bonds). The lipids are ordered in alphabetical order of the abbreviated name.

DG: diglyceride, LPC: lysophosphatidylcholine, LPE: lysophosphatidylethanolamine, PC: phosphatidylcholine, PE: phosphatidylethanolamine, SM: sphingomyelin, TG: triglyceride

### Supplementary Figures

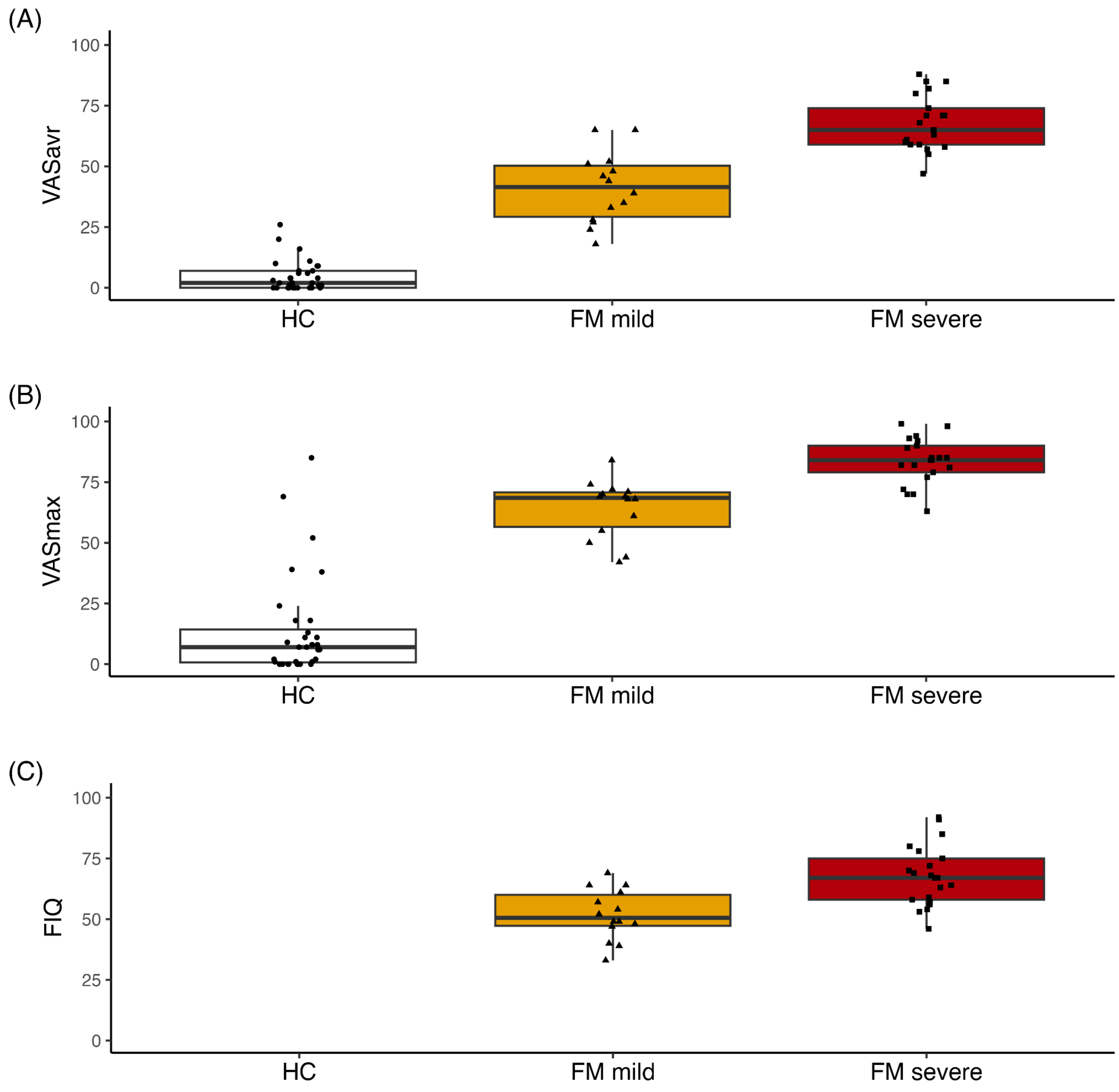


**Supplementary Figure S1. K-means clustering of the fibromyalgia (FM) subjects based on pain intensity ratings and disease severity.** The FM subjects were divided into two clusters, named FM mild and FM severe, through k-means clustering. The clustering was based on ratings of average and maximum pain on a Visual Analogue Scale (VAS) (VASavr and VASmax, respectively), as well as a fibromyalgia impact questionnaire (FIQ) score, representing disease severity. The resulting clusters were named after symptom severity. The FM subjects with more severe symptoms (FM severe) have higher VASavr (A), VASmax (B), and FIQ (C) scores than the FM subjects with mild symptoms (FM mild). The healthy control (HC) subjects were added to the plot to visualize the elevated pain intensity ratings.

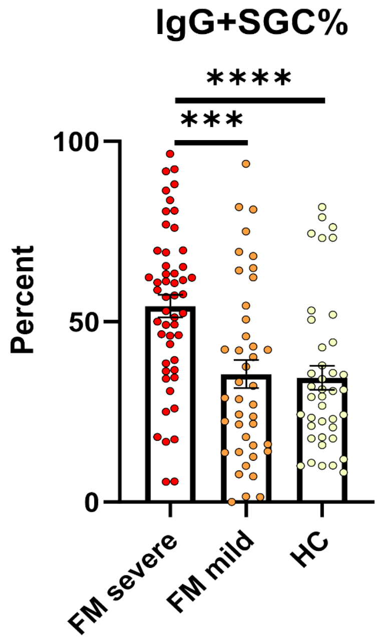


**Supplementary Figure S2. Percentage of IgG bound to satellite glial cells (SGC) for the FM patients and HC from the full cohort.** The groups were compared with a Kruskal-Wallis test followed by Dunn's post hoc test.

****: P < 0.0001, *** P < 0.001

**Supplementary Figures S3-S9.** Visualization of relative concentrations ($log_2$ intensity) of one lipid per lipid class. The boxplots illustrate differences between FM subjects and HC or FM severe, FM mild, and HC.

*ns: $P \geq 0.05$, *: $P < 0.05$, **: $P < 0.01$, ***: $P < 0.001$*

*HC: healthy controls, FM: subjects with fibromyalgia, FM mild: fibromyalgia subjects with mild symptoms, FM severe: fibromyalgia subjects with severe symptoms*

*DG: diglyceride, LPC: lysophosphatidylcholine, PC: phosphocholine, SM: sphingomyelin, TG: triglyceride.*

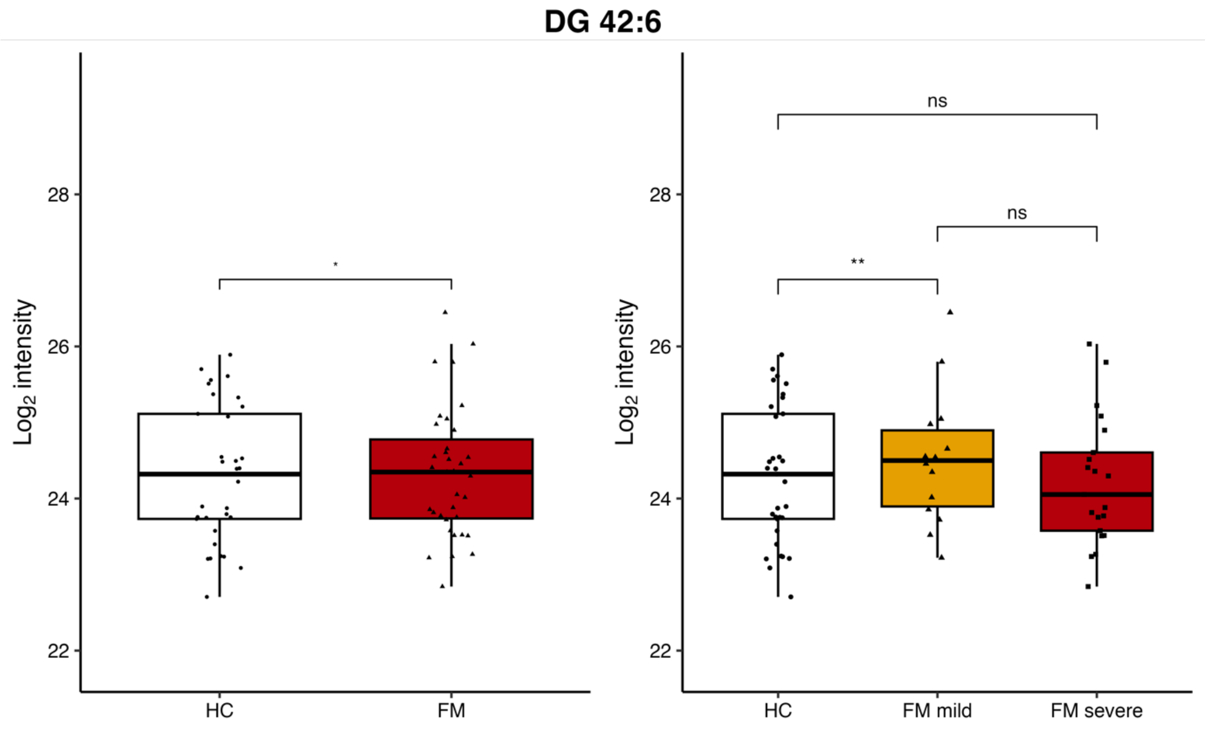


**Supplementary Figure S3. DG 20:3/22:3/0:0 (42:6).**

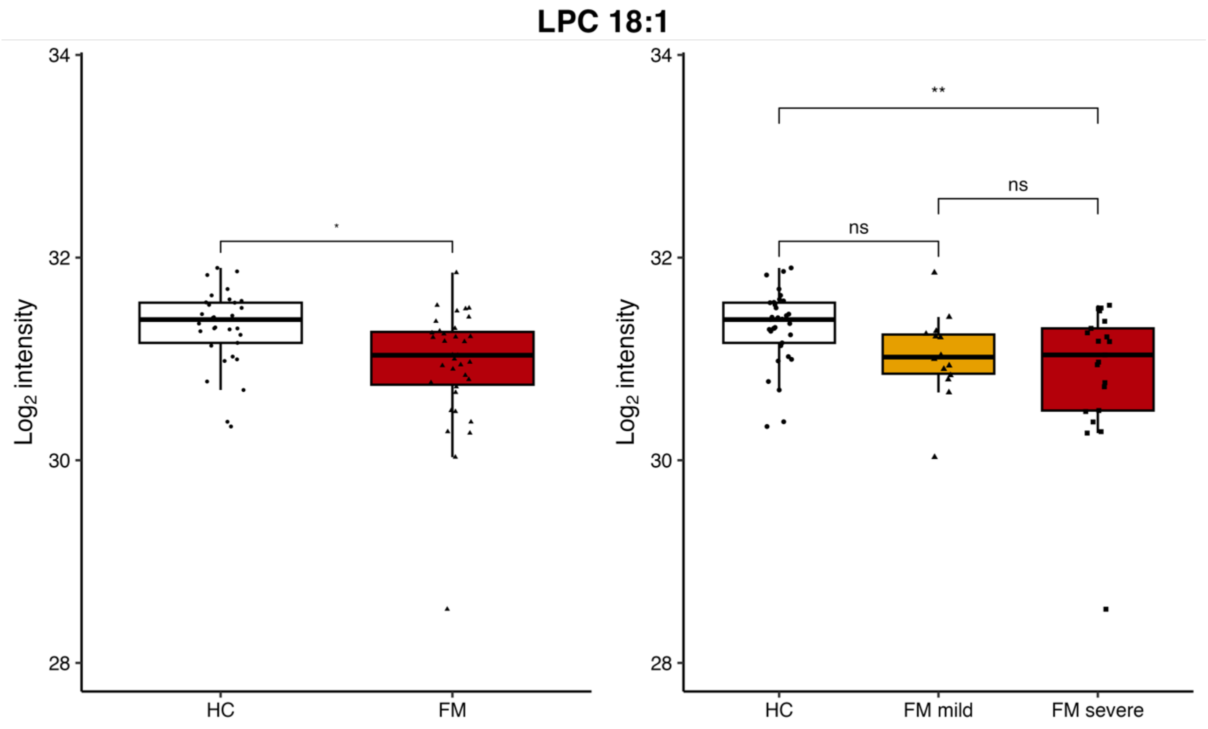


**Supplementary Figure S4. PC (18:1/0:0) (LPC 18:1).**

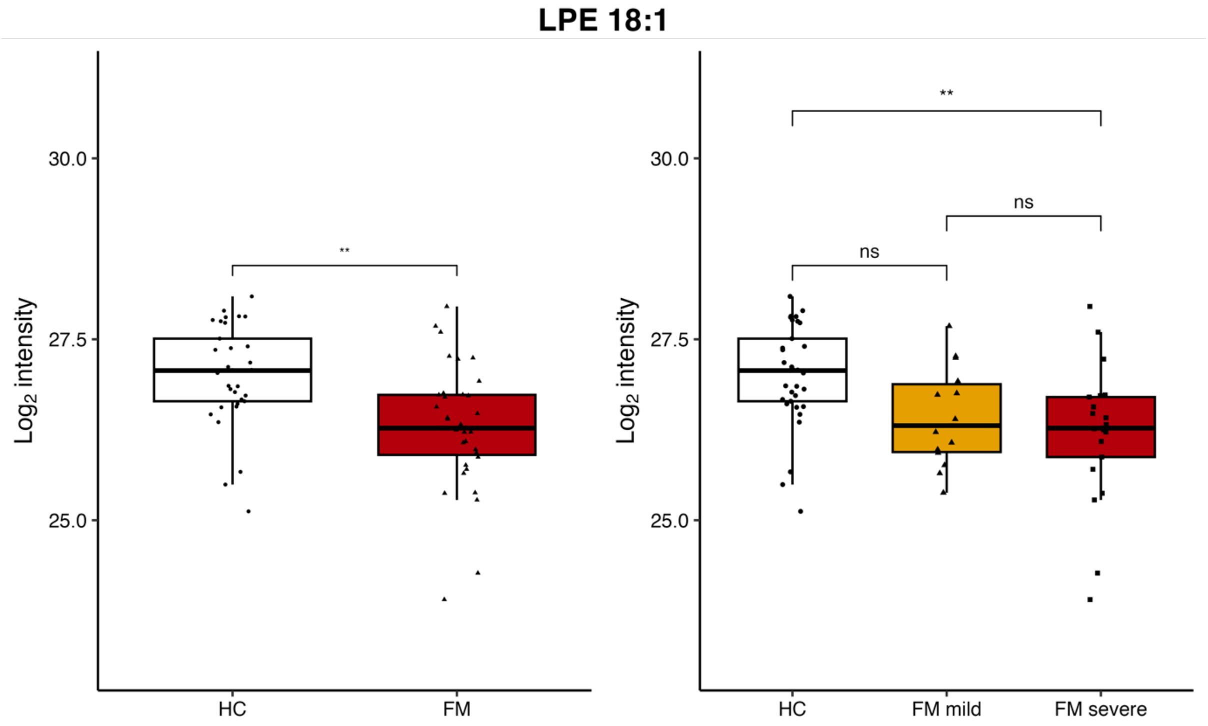


**Supplementary Figure S5. PE 18:1/0:0 (LPE 18:1).**

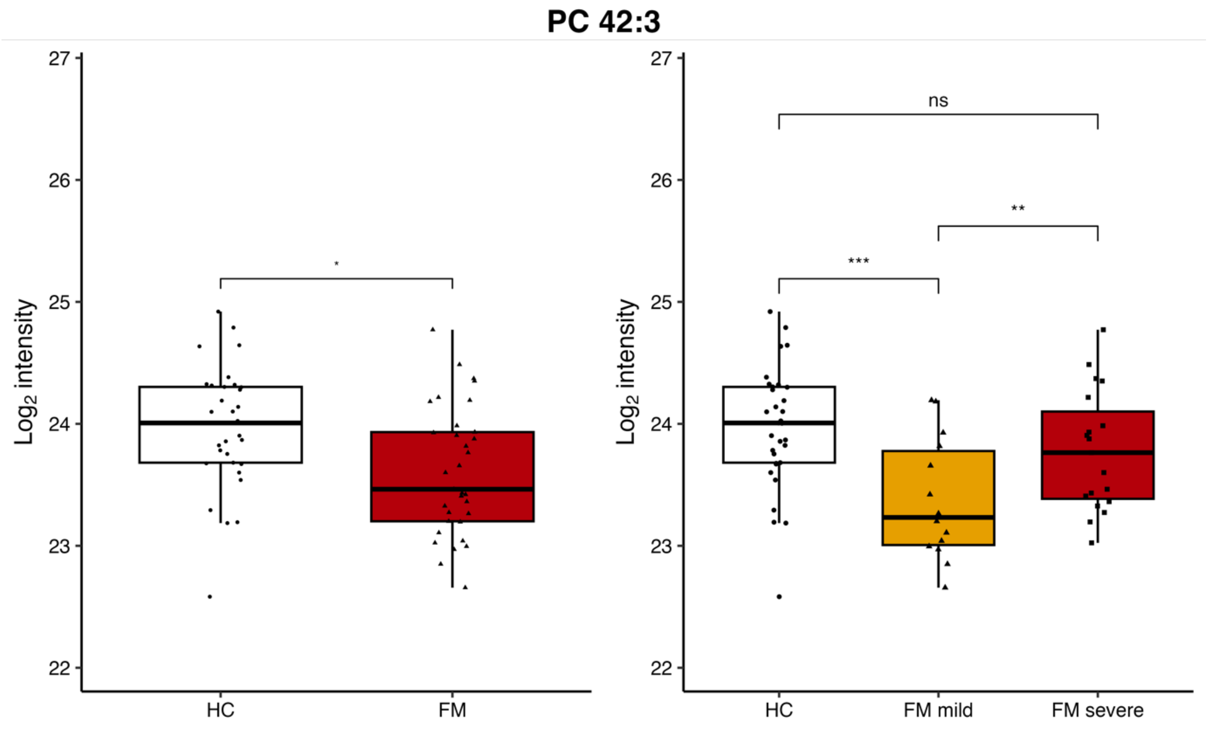


**Supplementary Figure S6. PC 18:3/24:0 (42:3).**

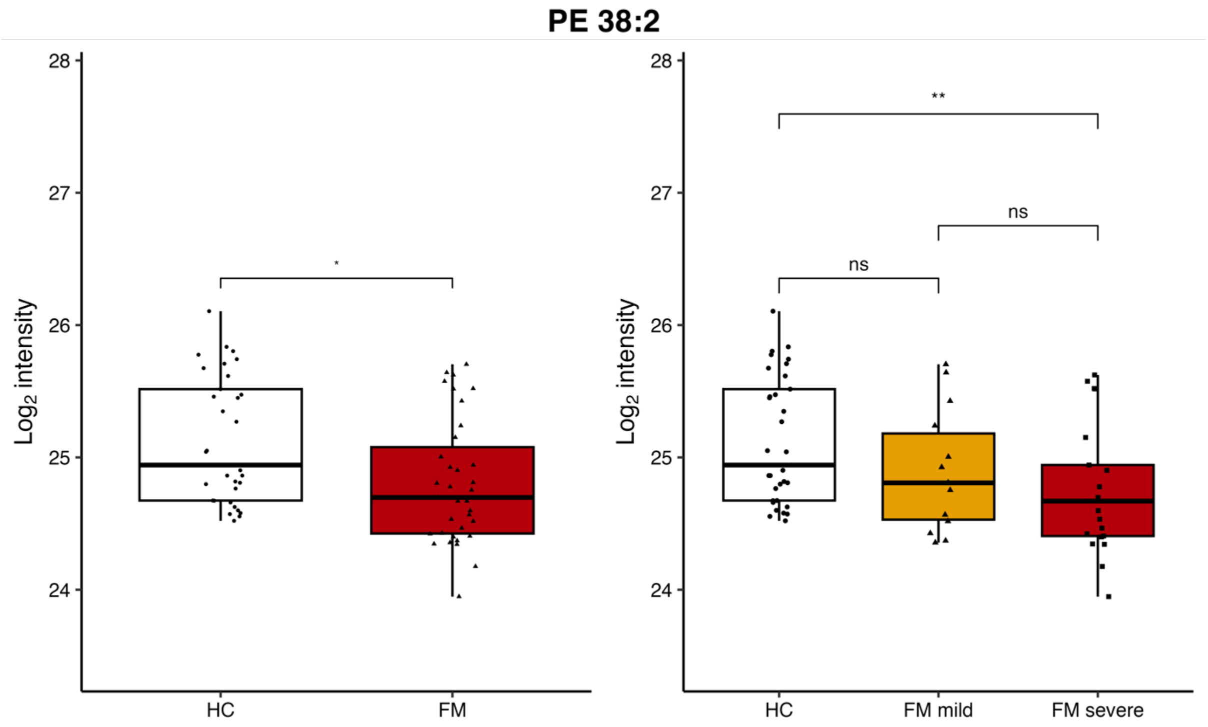


**Supplementary Figure S7. PE P-18:0/20:1 (38:2).**

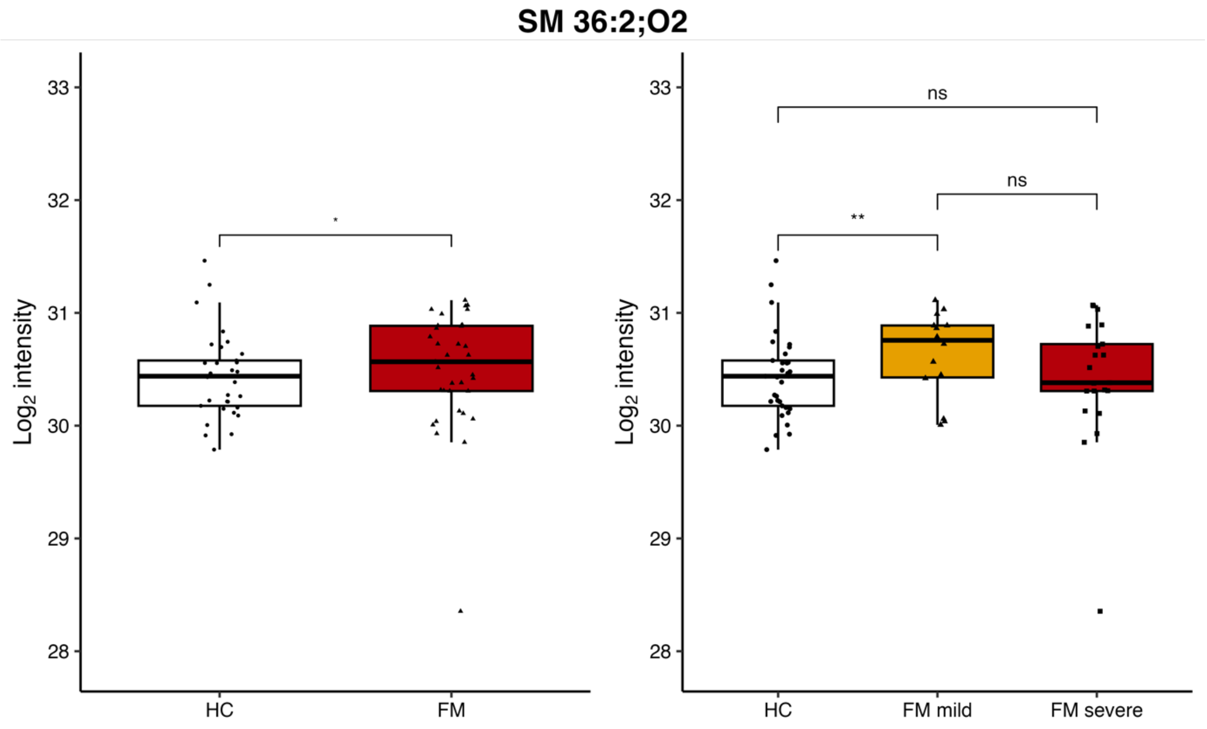


**Supplementary Figure S8. SM d18:1/18:1 (36:2;O2).**

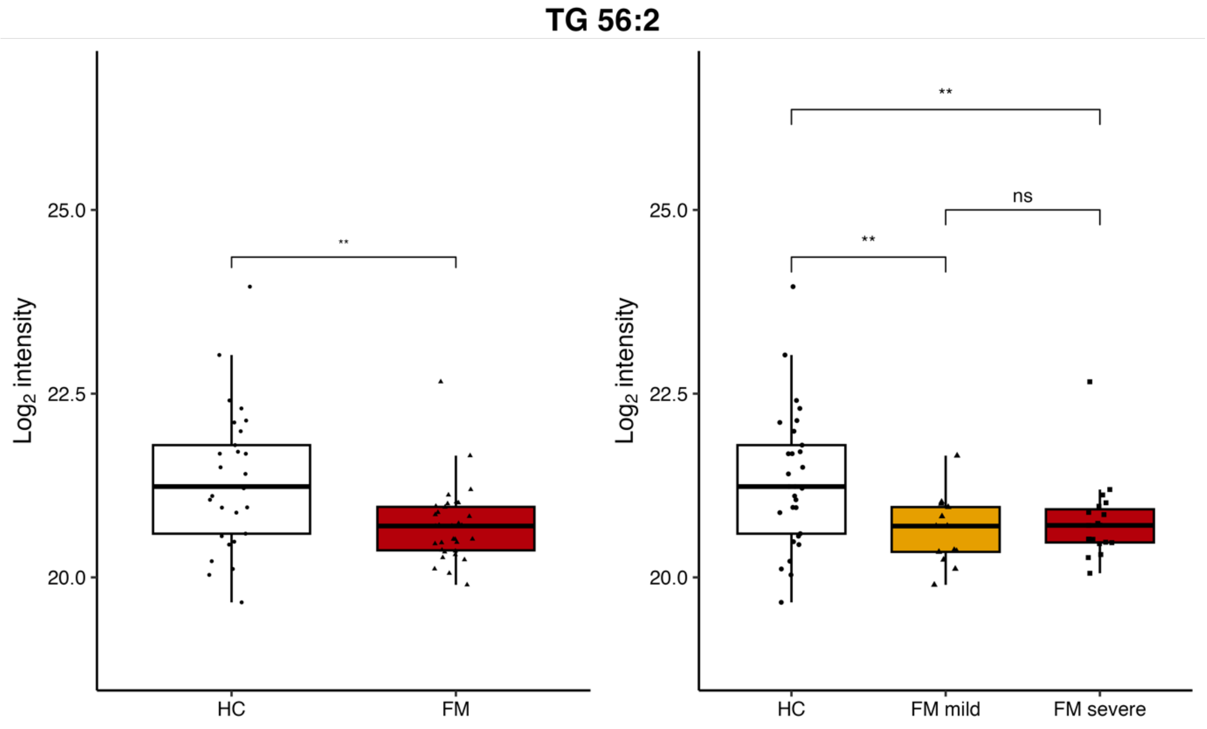


**Supplementary Figure S9. TG 12:0/22:1/22:1 (56:2).**
